# Retinoic acid-responsive *hox* genes in *hoxba* and *hoxbb* clusters direct pharyngeal pouch formation in zebrafish

**DOI:** 10.1101/2025.07.23.666262

**Authors:** Sohju Toyama, Nanami Hamano, Junpei Imagawa, Takumi Sugawara, Renka Fujii, Morimichi Kikuci, Yuki Kawabe, Akinori Kawamura

## Abstract

The segmented pharyngeal apparatus is crucial for organ development specific to vertebrates, and its formation relies on the proper development of pharyngeal pouches. While retinoic acid (RA) is known to influence pouch formation, the downstream genes involved have been unclear. In this study, we demonstrate that zebrafish mutants lacking both the *hoxba* and *hoxbb* clusters—teleost-specific duplicates of the ancestral *HoxB* cluster—exhibit a significant loss of posterior pharyngeal pouches and related skeletal elements. This phenotype resembles that observed in *raldh2* and *pax1a;pax1b* mutants. We identify *hoxb1a* and *hoxb1b* as RA-dependent genes expressed in the pharyngeal region that are essential for pouch formation. Morpholino-mediated knockdown of these genes replicated the pouch defects and decreased *pax1a* expression, indicating a regulatory pathway linking RA, *Hox*, and *pax1*. Our findings uncover a previously unrecognized role of *Hox* genes in early pouch segmentation and suggest that RA-responsive *HoxB* clusters were co-opted during vertebrate evolution to initiate pharyngeal regionalization.

## Introduction

During vertebrate embryogenesis, the pharyngeal region develops as a transiently segmented structure along the anterior–posterior axis, representing a unique morphological innovation acquired during vertebrate evolution (Graham and Smith, 2001; Kuratani, 2004). This region comprises pharyngeal arches—formed by migrating cranial neural crest cells and mesodermal cells—flanked externally by ectoderm-derived pharyngeal clefts and internally by endoderm-derived pharyngeal pouches, which are lateral outpocketings of the pharyngeal endoderm that separate adjacent arches. Through intricate interactons between these components, segmental identity is progressively established along the anterior-posterior axis. Each segment subsequently contributes to the formation of specific craniofacial structures and associated organs.

The segmented pharyngeal apparatus is conserved across cyclostomes, cartilaginous fishes, ray-finned fishes, and tetrapods, facilitating lineage-specific organogenesis (Graham and Smith, 2001; Kuratani, 2004). For example, in cyclostomes, pharyngeal arches form gill-supporting cartilages and branchial muscles; in fishes, they give rise to jaws, gill skeletons, and arterial arches; and in tetrapods, they contribute to the middle ear ossicles, pharyngeal musculature, and skeletal elements, as well as endoderm-derived organs such as the thyroid and thymus, supporting terrestrial adaptations including hearing and air breathing.

The formation of this segmented architecture critically depends on endoderm-derived pharyngeal pouch development. In zebrafish *van gogh* mutants, which lack pharyngeal endoderm, pharyngeal pouches fail to form, resulting in the complete absence of pharyngeal arches (Piotrowski and Nusslein-Volhard, 2000). Retinoic acid (RA) is also essential for pouch formation: in zebrafish and mouse mutants lacking *raldh2*, a key enzyme in RA biosynthesis, posterior pouches do not form, leading to the loss of corresponding arches (Begemann et al., 2001; Kopinke et al., 2006; Niederreither et al., 2003). Similarly, quail embryos subjected to vitamin A deficiency exhibit loss of posterior pouches due to impaired RA signaling (Quinlan et al., 2002). These findings demonstrate that diffusible RA is essential for pharyngeal pouch formation, suggesting a conserved mechanism among vertebrates.

Despite its critical role, the specific downstream genes regulated by RA signaling and the molecular mechanisms by which RA promotes pouch formation remain largely unknown. *Hox* genes are well-established downstream targets of RA in vertebrate embryogenesis and play pivotal roles such as axial patterning and neural tube patterning (Marshall et al., 1996). *Hox* genes are arranged in clusters, and their linear genomic organization corresponds to spatial and temporal expression domains. While mice and humans possess 39 *Hox* genes across four clusters (*HoxA–D*) following two rounds of whole-genome duplication, teleost fishes such as zebrafish underwent an additional duplication, resulting in seven clusters comprising 49 *hox* genes (Amores et al., 1998). Although many *Hox* gene knockout mice, including their compound mutants, have been generated, none exhibit a complete loss of posterior pharyngeal pouches comparable to that observed in RA-deficient mutants, leaving the contribution of *Hox* genes to pouch formation unresolved. In our previous study, we generated a complete series of zebrafish mutants in which each of the seven *hox* clusters was individually deleted (Yamada et al., 2021). In this study, we show that mutants lacking both *hoxba* and *hoxbb* clusters—teleost-specific duplicates of the ancestral *HoxB* cluster—exhibited a marked loss of jaw skeletal elements. This phenotype is attributed to defects in posterior pharyngeal pouch formation and was not observed in other combinations of zebrafish *hox* cluster mutants. Further analysis reveals that *hoxb1a* and *hoxb1b*, two paralogous genes, are transcriptionally regulated by RA and are required for pharyngeal pouch formation. Our findings support a model in which RA signaling initiates segmentation of the pharyngeal endoderm via activation of *Hox* genes, particularly within *HoxB*-related clusters. This developmental axis likely contributed to the emergence of pharyngeal structures across vertebrate lineages.

## Results

### Severe defects in pharyngeal skeleton formation in zebrafish *hoxba;hoxbb* cluster mutants

In a previous study, we generated seven zebrafish mutants, each deficient in a single *hox* cluster (Yamada et al., 2021). Examination of the pharyngeal skeleton cartilage in these mutants revealed that only homozygous *hoxba* cluster mutants (hereafter referred to as *hoxba* mutants) exhibited abnormalities in jaw cartilage formation, specifically altered morphology of the ceratohyals (Yamada et al., 2021). However, these malformations were relatively mild and did not result in the loss of any pharyngeal skeleton components other than the basihyal (Figure 1B), suggesting functional compensation by other *hox* clusters. A likely candidate for such compensation is *hoxbb* cluster, which, along with *hoxba* cluster, originated from the ancient *HoxB* cluster via the teleost-specific whole-genome duplication (Amores et al., 1998). Supporting this hypothesis, our previous work demonstrated cooperative roles of *hoxba* and *hoxbb* clusters in early pectoral fin bud development in zebrafish (Kikuchi et al., 2025). To assess their functional redundancy in pharyngeal development, we intercrossed *hoxba^+/-^*;*hoxbb^+/-^* mutants and analyzed the jaw cartilage in larvae at 5 dpf using Alcian blue staining. In wild-type pharyngeal skeletons, the pharyngela skeleton includes a midline basibranchial and five pairs of ceratobranchials aligned along the anterior-posterior axis (Figure 1A). In *hoxba;hoxbb* double mutants, only the anterior portion of the basibranchial and the first pair of ceratobranchials were present, both with altered morphologies, while the remaining basibranchial and the second to fifth ceratobranchials were absent (Figure 1D). The ceratohyals were even more deformed, showing reversed orientation. Compared to *hoxba^-/-^* or *hoxbb^-/-^* single mutants (Figure 1B, C), *hoxba;hoxbb* double mutants displayed markedly more severe morphological defects in the jaw skeleton, suggesting that *hoxba* and *hoxbb* clusters function redundantly in pharyngeal skeleton formation.

**Figure 1.**
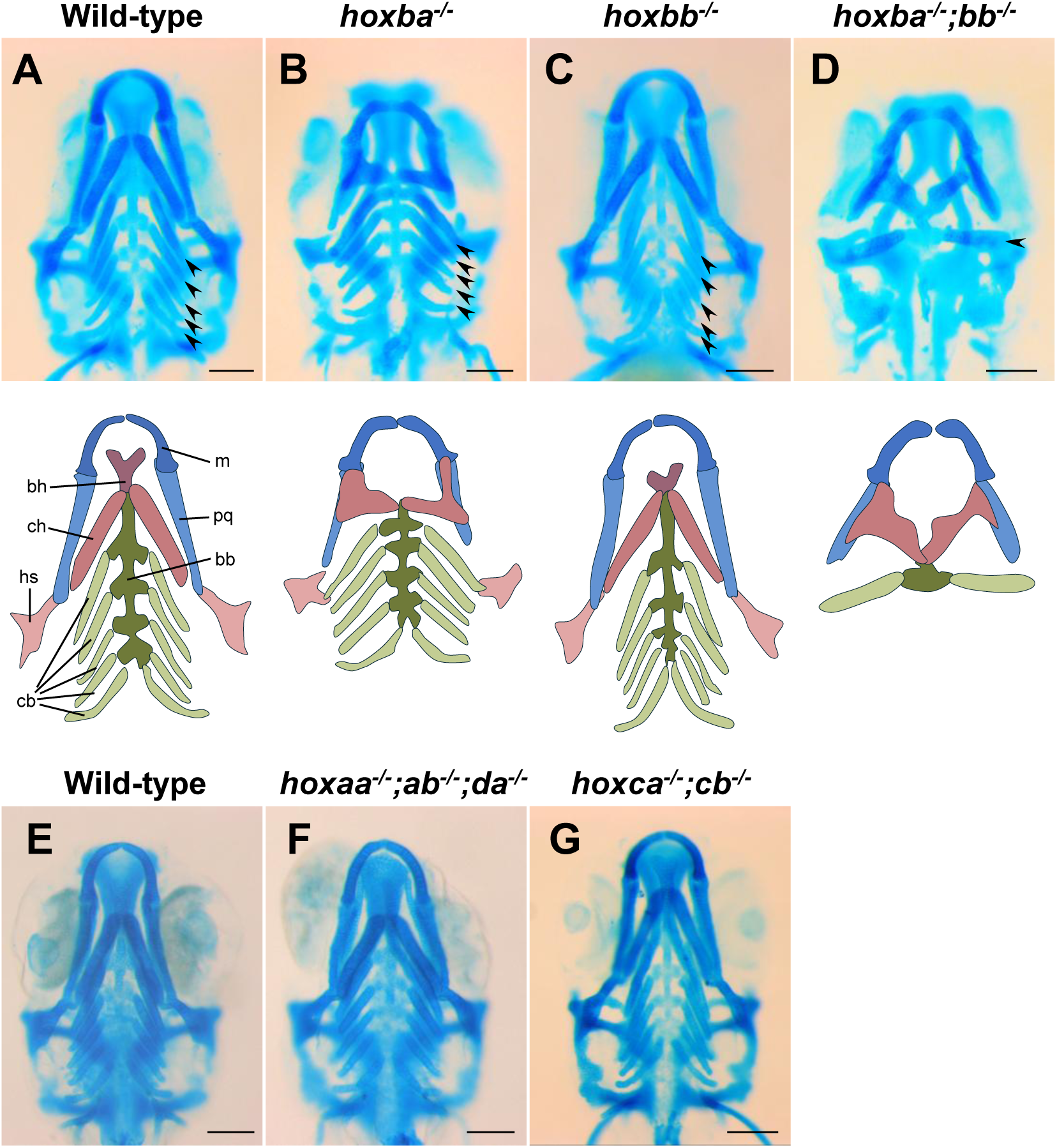
Malformations of jaw cartilage in zebrafish with deleted *hoxba;hoxbb* clusters. (A-D) The upper panels show ventral views of the jaw cartilages from 5 dpf Aclian blue-stained embryos, resulting from intercrosses between *hoxba;hoxbb* hemizygous fish. The lower panels provide schematic diagrams illustrating the skeletal components of the ventral jaws: m; Meckel’s cartilage, bh; basihyal, ch; ceratohyal, hs; hyosympletic, pq; palatoquadrate, bb; basibranchial, cb; ceratobranchial (indicated by arrowheads). (E-G) No apparent morphological abnormalities of jaw cartilages were observed in *hoxaa;hoxab;hoxda* triple and *hoxca;hoxcb* double homozygous mutants at 5dpf. As no morphological abnormalities were noted, a schematic diagram is not provided. For each genotype, at least three samples were examined to confirm reproducibility. Scale bar: 100 μm.

Taking advantage of our availability of seven individual *hox* cluster mutants in zebrafish, we next investigated whether similarly severe jaw skeletal defects occur in other combinations of *hox* cluster deficiencies. In mice, *HoxA* and *HoxD* clusters have been shown to function redundantly in limb development (Kmita et al., 2005). Similarly, in zebrafish, overlapping functions among *hoxaa*, *hoxab*, and *hoxda* clusters—homologous to the mouse *HoxA* and *HoxD*—have been implicated in pectoral fin development (Ishizaka et al., 2024). Accordingly, we generated zebrafish triple homozygous mutants for *hoxaa*, *hoxab*, and *hoxda* clusters and analyzed the pharyngeal skeletons at 5 dpf. However, no significant morphological abnormalities were observed in their jaw skeletons compared to sibling wild-types (Figure 1E, F). Similarly, *hoxca;hoxcb* double mutants displayed no appreciable defects in pharyngeal skeleton morphology (Figure 1G). These findings indicate that *hoxba* and *hoxbb* clusters— both derived from the ancient *HoxB* cluster—play a predominant role in the formation of jaw structures in zebrafish.

### Segmental formation of posterior pharyngeal pouches is severely impaired in zebrafish *hoxba;hoxbb* mutants

In *hoxba;hoxbb* double mutants, most components of the pharyngeal skeleton-including the basibranchials and ceratobranchials except for the anteriormost ones, are present. This phenotype resembles that of *raldh2* (*retinaldehyde dehydrogenase 2*) mutants, which are deficient in RA synthesis (Begemann et al., 2001; Grandel et al., 2002), and also resembles that of *pax1a;pax1b* double mutants in zebrafish and medaka (Liu et al., 2020; Okada et al., 2016; Okada and Takada, 2020). In both cases, only the first and second pair of pharyngeal pouches, derived from the pharyngeal endoderm, form, while the third and more posterior pharyngeal pouches fail to develop. To determine whether *hoxba;hoxbb* mutants show a similar defect, we performed immunostaining with the zn-8 antibody. In wild-type embryos at 30 hpf, five pharyngeal pouches were observed at regular intervals along the anterior-posterior axis (Figure 2A, A’). While no obvious abnormalities were seen in *hoxba* or *hoxbb* single mutants, *hoxba;hoxbb* double mutants displayed a phenotype similar to that of *raldh2* mutants: absence of the third and more posterior pharyngeal pouches (Figure 2B-E, B’-E’). To further assess this defect, we examined the expression patterns of *pax1a* by whole-mount *in situ* hybridization (Figure 2F-J). In wild-type, *hoxba* mutants, or *hoxbb* single mutants, *pax1a* was expressed periodically in five pharyngeal pouches (Figure 2F-H, F’-H’, F’’-H’’). In contrast, *hoxba;hoxbb* double mutants showed markedly reduced *pax1a* expression in the second and subsequent pharyngeal pouches, similar to that of *raldh2* mutants (Figure 2I-J, I’-J’, I’’-J’’), consistent with zn-8 immunostaining results. These findings reveal a previously unrecognized role for *hox* genes: *hoxba* and *hoxbb* clusters cooperate in the segmental formation of pharyngeal pouches.

**Figure 2.**
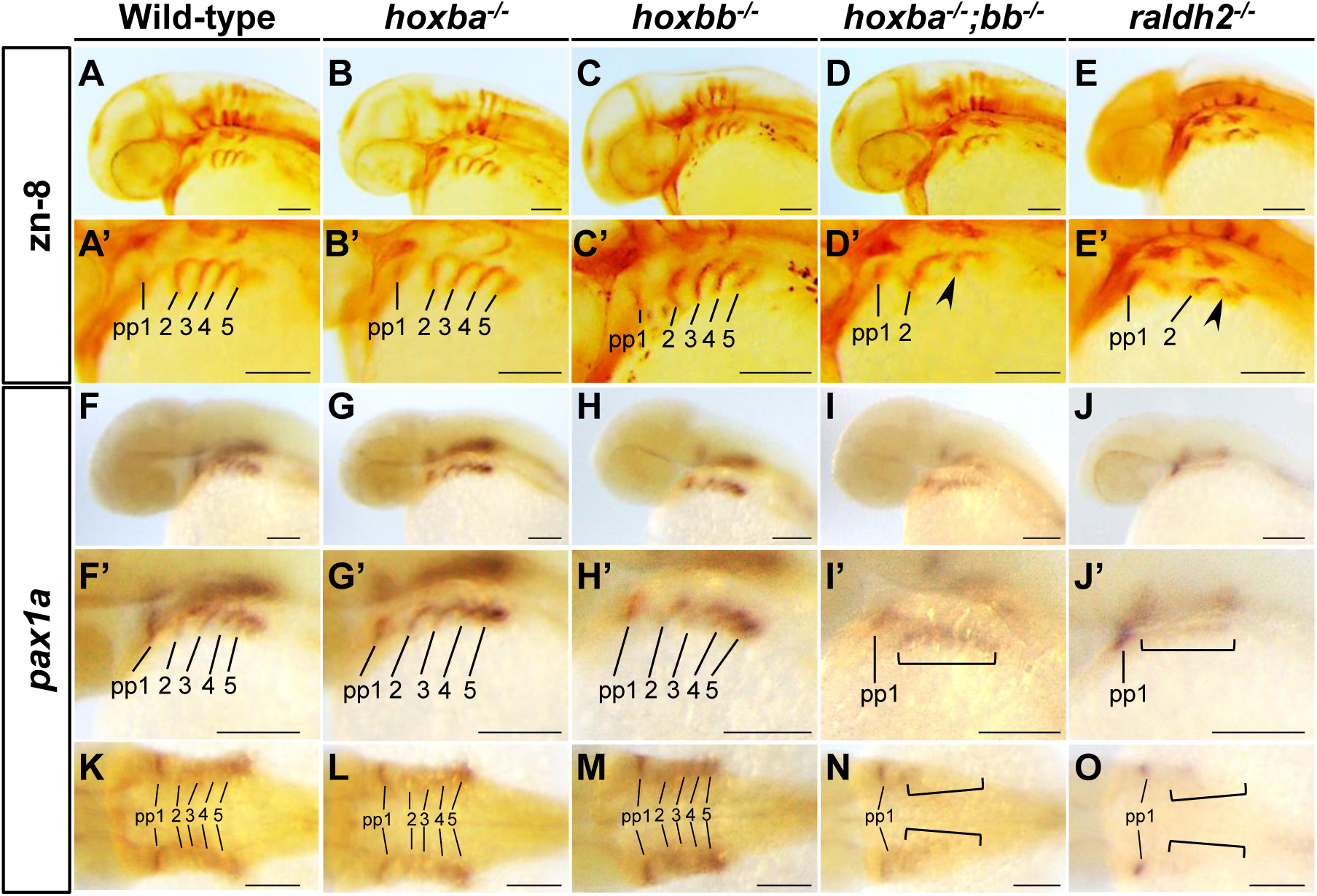
Zebrafish *hoxba;hoxbb* mutants exhibit deformation of the posterior pharyngeal pouches. (A-E) The upper panels display oblique images of 36 hpf embryos stained with the zn-8 monoclonal antibody. Embryos in panels A-D were obtained from intercrosses between *hoxba;hoxbb* hemizygous fish. Panel E shows an immunostained *raldh2* homozygous embryo at 36 hpf. The lower panels A’-E’ present magnified images of the pharyngeal region from the upper panels, with pharyngeal pouches 1-5 indicated. The arrowheads in panels D’ and E’ denote severely disorganized and partially absent lateral pharyngeal endoderm. (F-J) Expression patterns of *pax1a* in 36 hpf embryos were analyzed using whole-mount *in situ* hybridization. The upper panels F-J present oblique images of 36 hpf embryos exhibiting *pax1a* expression. The lower panels F’-J’ show magnified images of the pharyngeal region from the upper panels, with pharyngeal pouches 1-5 indicated. Presumptive posterior pharyngeal pouches with significantly reduced expression of *pax1a* are emphasized by brackets in panels I’ and J’. Images F’’-J’’ were taken from the dorsal side of the same stained embryo shown in F-J. For each genotype, at least three samples were examined to confirm reproducibility. Scale bar: 100 μm.

### Expression of *hoxb1a* and *hoxb1b* in the pharyngeal regions is RA-dependent

We next sought to identify specific *hox* genes within *hoxba* and *hoxbb* clusters responsible for pharyngeal pouch formation. The severe phenotype observed only in *hoxba;hoxbb* double mutants suggests the involvement of duplicated paralogous *hox* genes derived from the ancestral *HoxB* cluster. Furthermore, the similarity to *raldh2* mutants suggests that these *hox* genes act as downstream targets of RA signaling, a known regulator of vertebrate development (Boncinelli et al., 1991; Langston and Gudas, 1994; Marshall et al., 1996). We focused on *hoxb1a* and *hoxb1b*, known for their roles in hindbrain patterning (McClintock et al., 2002; Weicksel et al., 2014), though their function in pharyngeal pouch development is unreported. At the 10-12-somite stage, *hoxb1a* expression was weakly detected in the pharyngeal region-likely including the precursor cells of the pharyngeal pouches in sibling embryos (Figure 3A, A’, E). This expression was markedly reduced in *raldh2* mutants, while expression in the rhombomere remained unchanged (Figure 3B, B’). At 36 dpf, when the pharyngeal arches are established, the expression of *hoxb1a* in the pharyngeal region was almost undetectable in *raldh2* mutants (Figure 3E, F). Likewise, *hoxb1b* expression was observed in the pharyngeal region at both stages and also significantly reduced in *raldh2* mutants (Figure 3C, D, C’, D’, G, H, G’, H’). These data indicate that *hoxb1a* and *hoxb1b* are expressed in RA-dependent manner in the pharyngeal region and are strong candidates for mediating pharyngeal pouch formation.

**Figure 3.**
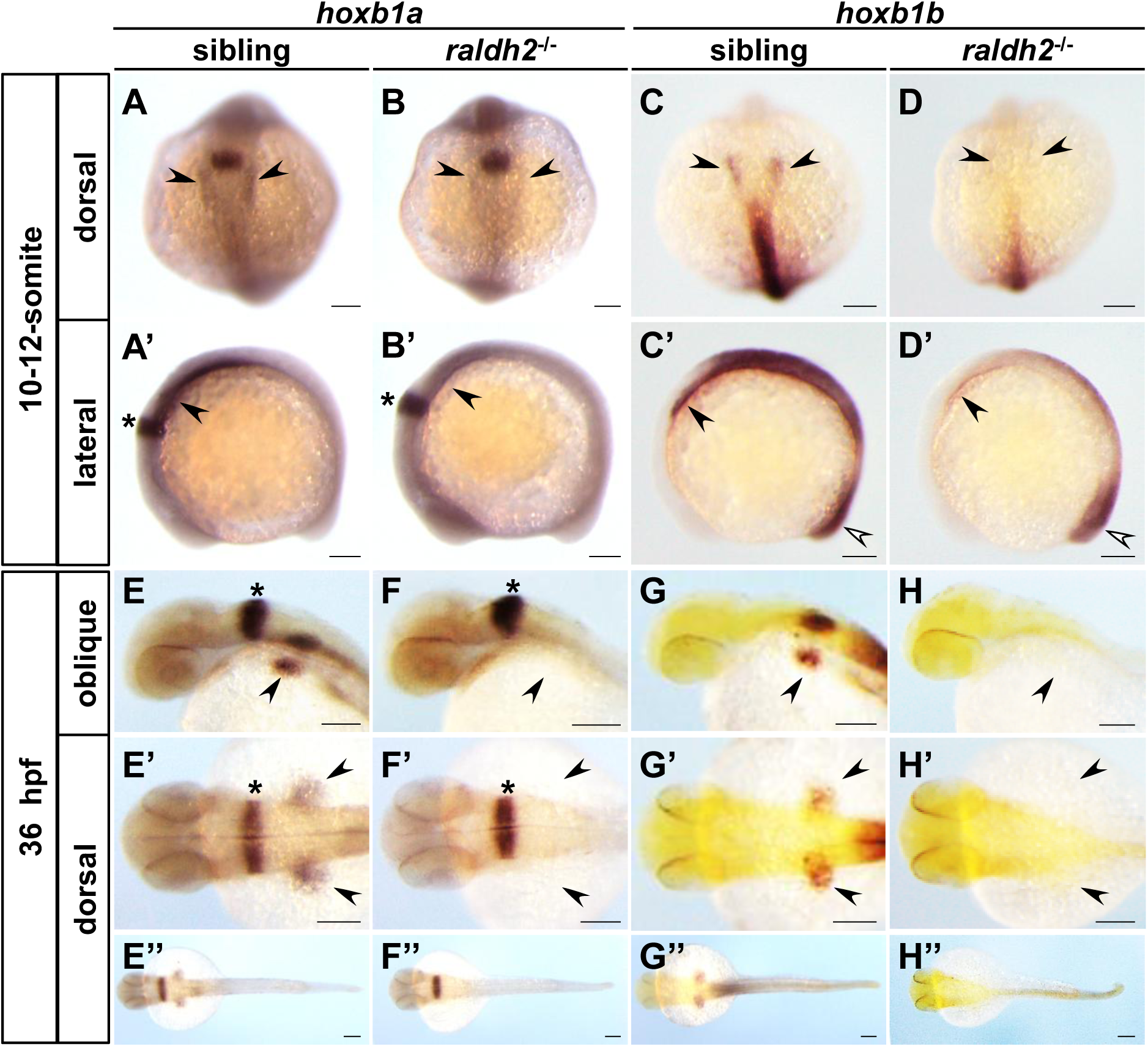
Expression of *hoxb1a* and *hoxb1b* in the pharyngeal regions depends on RA. Expression patterns of zebrafish *hoxb1a* and *hoxb1b* were analyzed in sibling and *raldh2* homozygous embryos using whole-mount *in situ* hybridization. Embryos at the 10- to 12-somite stages and at 36 hpf were examined. The images of the same specimens were taken from the dorsal (A-D) and lateral views (A’-D’). Black arrowheads in panels A-H indicate the positions of presumptive pharyngeal regions, including pharyngeal pouches. White arrowheads in panels C’ and D’ denote the expression of *hoxb1b* in the tailbud. Asterisks represent the specific expression of *hoxb1a* in rhombomere 4. For each result, at least three samples were analyzed to confirm reproducibility. Scale bar: 100 μm.

### Knockdown of *hoxb1a* and *hoxb1b* results in severe pharyngeal pouch malformations similar to *hoxba;hoxbb* mutants

To investigate whether *hoxb1a* and *hoxb1b* contribute to pharyngeal pouch formation, we analyzed the phenotypes resulting from the loss of function of these genes. Frameshift mutants of zebrafish *hoxb1a* and *hoxb1b* have already been generated with morphological abnormalities in the hindbrain reported in both *hoxb1b* single mutants and *hoxb1a;hoxb1b* double mutants (Weicksel et al., 2014). However, defects in pharyngeal pouch formation have not been described. In our previous study, we observed that the pectoral fin loss phenotype seen in *hoxba;hoxbb* cluster deletion mutants was not fully recapitulated in frameshift mutants of candidate *hox* genes, with the phenotype being markedly milder (Kikuchi et al., 2025). This discrepancy may be explained by a mechanism known as transcriptional adaptation, in which mRNAs containing premature stop codons induced by frameshift mutations trigger the upregulation of structurally similar transcripts, thereby compensating for the loss of function (El-Brolosy et al., 2019; Ma et al., 2019). To avoid the confounding effects of transcriptional adaptation, we performed functional knockdown experiments using antisense morpholino oligos previously validated to inhibit these genes (McClintock et al., 2002), which do not induce transcriptional adaptation. Injection of morpholinos targeting either *hoxb1a* or *hoxb1b* alone, together with a negative control morpholino, did not result in noticeable abnormalities (Table S1). However, combined knockdown of both *hoxb1a* and *hoxb1b* caused severe defects in pharyngeal pouch formation, resembling *hoxba;hoxbb* mutant phenotype (Figure 4A, A’, B, B’). Given the significant reduction of *pax1a* expression in *hoxba;hoxbb* mutants, we also examined *pax1a* expression in embryos deficient in *hoxb1a* and *hoxb1b*. While the first pharyngeal pouch remained unaffected, *pax1a* expression in the second and more posterior pouches was markedly reduced (Figure 4C, C’, D, D’). These results suggest that *hoxb1a* and *hoxb1b*, regulated by RA, cooperatively control *pax1a* expression during zebrafish pharyngeal pouch formation. Collectively, these findings provide, to our knowledge, the first evidence for a novel role of *Hox* genes in the pharyngeal pouch development in vertebrates.

**Figure 4.**
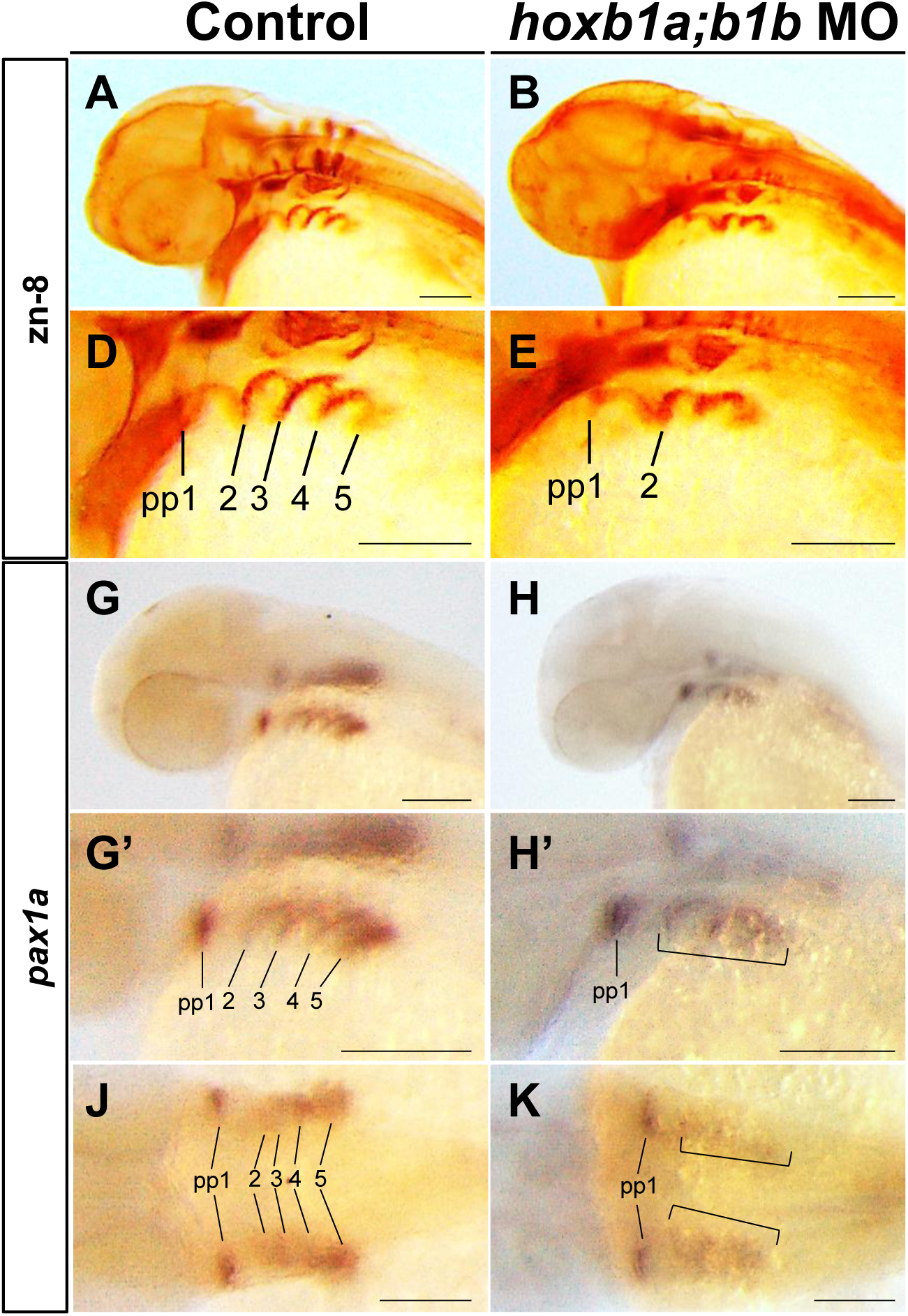
*hoxb1a* and *hoxb1b* morpholino-injected embryos exhibit severe deformation of pharyngeal pouches. (A, B) Panels A and B display oblique images of morpholino-injected embryos at 36 hpf, stained with the zn-8 monoclonal antibody. As a control, negative control morpholino oligos were injected. Panels A’ and B’ present magnified images of the pharyngeal region from panels A and B, with pharyngeal pouches 1-5 indicated. The arrowhead in panel B’ denotes severely disorganized and partially absent lateral pharyngeal endoderm. (C, D) Expression patterns of zebrafish *pax1a* were analyzed in morpholino-injected embryos at 36 hpf using whole-mount *in situ* hybridization, shown in oblique views (C, D), magnified images of the pharyngeal regions (C’, D’), and dorsal views (C’’, D’’). Presumptive posterior pharyngeal pouches with significantly reduced expression of *pax1a* are marked by brackets in panels D’ and D’’. For each result, at least three samples were analyzed to confirm reproducibility. Scale bar: 100 μm.

## Discussion

### Essential *Hox* genes in early pharyngeal pouch formation

Previous studies have shown that RA is a conserved regulator of posterior pharyngeal pouch formation across vertebrates (Begemann et al., 2001; Niederreither et al., 2003; Quinlan et al., 2002; Wendling et al., 2000). *Hox* genes are recognized as potential downstream effectors of RA signaling. In *Raldh2* knockout mice, which lack the enzyme required for RA synthesis, the expression of *Hoxa1* and *Hoxb1* in the pharyngeal pouches is significantly reduced (Niederreither et al., 2003; Wendling et al., 2000), suggesting their involvement in pouch formation. However, mice with single or double knockouts of *Hoxa1* and *Hoxb1*, show defects primarily in pharyngeal arch identity—mainly derived from cranial neural crest cells—but no clear abnormalities in pharyngeal pouch formation have been reported (Chisaka et al., 1992; Goddard et al., 1996; Rossel and Capecchi, 1999). Similarly, mice lacking most genes in *HoxB* cluster (excluding *Hoxb13*), homologous to the zebrafish *hoxba* and *hoxbb* clusters, do not display significant defects in pouch development (Medina-Martinez et al., 2000). While *Hoxa3* knockout mice exhibit thymic abnormalities originating from pharyngeal pouch derivatives (Manley and Capecchi, 1995), no single or compound *Hox* gene knockout model has recapitulated the extensive loss of posterior pharyngeal pouches observed in this study. These results suggest that prior loss-of-function studies in mice have provided only limited direct evidence for the role of *Hox* genes as downstream effectors of RA in early pouch formation. In this study, we present genetic evidence from zebrafish demonstrating that mutants lacking both the *hoxba* and *hoxbb* clusters—paralogous clusters derived from the ancestral *HoxB* cluster—exhibit severe loss of posterior pharyngeal pouches, resulting in the loss of jaw skeletal elements (Figure 1, 2), a phenotype similar to *raldh2* mutants with impaired RA synthesis (Begemann et al., 2001). Our findings suggest that *Hox* genes regulated by RA are essential for initiating the segmentation of through posterior pouch formation. Based on previous studies and this study, we propose a stepwise model in which initila segmentation by early acting *Hox* genes is followed by the establishment of anterior–posterior identity in each arch and pouch through later-acting *Hox* genes.

### Phenotypic discrepancy of *Hox* mutants between zebrafish and mice

We showed that zebrafish mutants of *hoxba;hoxbb* cluster and *hoxb1a;hoxb1b* knockdown embryos exhibit a marked loss of posterior pharyngeal pouches (Figure 2, 4). However, as noted above, comparable phenotypes have not been observed in mice lacking *HoxB* cluster (except for *Hoxb13*) or in *Hoxb1* knockout mice (Goddard et al., 1996; Medina-Martinez et al., 2000), highlighting a phenotypic discrepancy between homologous *Hox* genes in zebrafish and mice. A similar divergence is seen in the development of vertebrate paired appendages: zebrafish *hoxba;hoxbb* mutants lack pectoral fins (Kikuchi et al., 2025), whereas *HoxB* cluster-deficient mice do not exhibit homologous forelimb loss (Medina-Martinez et al., 2000). In mice, extensive functional redundancy among paralogous or adjacent *Hox* genes has been well demonstrated through genetic studies (Horan et al., 1995; Wellik and Capecchi, 2003; Wellik et al., 2002). In contrast, our recent studies in zebrafish and medaka have shown that *Hox* genes responsible for dorsal and anal fin development reside mainly within *HoxC*-related clusters (Adachi et al., 2024; Koita et al., 2025), whereas those essential for pectoral fin formation are specifically found in *HoxB*-related clusters (Kikuchi et al., 2025). In this study, we generated multiple hox cluster-deletion mutants and identified the *HoxB-*related cluster as critical for pharyngeal pouch development. These results suggest that in zebrafish, *Hox* genes responsible for specific developmental processes may be functionally concentrated within distinct *Hox* clusters, potentially a result of teleost-specific whole-genome duplication. This genomic architecture may have unveiled a previously unrecognized role for *Hox* genes, including their essential contribution to pharyngeal pouch formation. We presume that these functions in pharyngeal pouch formation are likely conserved in other vertebrates, including mice, but have remained undetected in murine models due to functional redundancy among *Hox* genes.

### Insights into the molecular mechanisms of pharyngeal pouch formation

This study demonstrates that *hoxb1*a and *hoxb1b*, functioning downstream of RA signaling, are essential for the early development of pharyngeal pouches in zebrafish (Figure 3, 4). Multiple retinoic acid response elements (RAREs) have been identified near the zebrafish *hoxb1a* and *hoxb1b* loci (Ishioka et al., 2011), suggesting that their expression in the pharyngeal region is regulated by RA. Similarly, RAREs are involved in the regulation of *Hoxb1* expression in mice (Huang et al., 2002; Marshall et al., 1994), supporting a conserved RA-RARE-Hox regulatory axis in pharyngeal pouch formation. Furthermore, zebrafish *hoxba;hoxbb* mutants and *hoxb1a;hoxb1b* knockdown embryos exhibit phenotypes resembling those of *pax1a;pax1b* double mutants, characterized by defective formation of posterior pharyngeal pouches (Okada et al., 2016; Okada and Takada, 2020). Notably, *pax1a* expression in the pharyngeal pouch is significantly reduced in these *hox*-deficient embryos, indicating that *pax1* functions downstream of *hox* genes activated by RA signaling and plays a role in the molecular pathways underlying pharyngeal pouch development.

### Evolutionary implications

The segmental structure of the pharyngeal apparatus, a vertebrate-specific innovation, contributes to the development of vertebrate-specific organs such as the jaw and thymus. As pharyngeal pouches play a central role in forming this structure, elucidating how vertebrates acquired these structures is a major question in evolutionary developmental biology. Our findings show that *Hox* genes are indispensable for posterior pharyngeal pouch formation in zebrafish. *Hox* genes encode evolutionarily conserved transcription factors found across bilaterian animals, including species that lack pharyngeal structures. This suggests that certain molecular changes in *Hox* genes— particularly the acquisition of RA-dependent regulatory mechanisms in the pharyngeal regions—may have enabled *Hox* genes to be co-opted into pharyngeal pouch formation during vertebrate evolution. Thus, this study not only reveals the novel role of *Hox* genes in pharyngeal pouch development but also provides critical insights into the molecular mechanisms underlying the evolutionary acquisition of these vertebrate-specific structures. Understanding the precise molecular changes that endowed *Hox* genes with this new role will be crucial for uncovering the genetic basis of organ innovation in early vertebrate evolution.

### Limitation of this study

In this study, we investigate the roles of *hoxb1a* and *hoxb1b* as candidate *hox* genes in the formation of pharyngeal pouches. In early zebrafish embryos, the pharyngeal region is not yet clearly defined, and both *hoxb1a* and *hoxb1b* are expressed throughout the pharyngeal regions, including the pharyngeal arches and pouches. This widespread expression complicates the distinction between these structures. While using double mutants that lack both *hoxb1a* and *hoxb1b* would be a suitable approach to assess their contributions to pharyngeal pouch formation, these mutants may exhibit transcriptional adaptation, which could complicate the interpretation of results. To address this, we employed antisense morpholino-mediated knockdown to explore the functional roles of these genes in pharyngeal pouch development.

### Experimental model

#### Zebrafish lines and maintenance

Zebrafish (Riken WT; RW) were provided by the National BioResource Project (NBRP) Zebrafish in Japan. Zebrafish were maintained at 27°C with a 14-h light/10-h dark cycle. Embryos were obtained from natural spawning and raised at 28.5°C. Developmental stages of the embryos were determined based on hours post-fertilization (hpf) or days post-fertilization (dpf) and morphological features as previously described (Kimmel et al., 1995). Zebrafish mutants used in this study are *hoxaa* cluster-deficient*^sud111^*, *hoxab* cluster-deficient*^sud112^*, *hoxba* cluster-deficient*^sud113^*, *hoxbb* cluster-deficient*^sud114^*, *hoxca* cluster-deficient*^sud115^*, *hoxcb* cluster-deficient*^sud124^*, *hoxda* cluster-deficient*^sud116^*, and *raldh2^sud118^*, which were all generated by using the CRISPR-Cas9 system in our previous studies (Kikuchi et al., 2025; Yamada et al., 2021). All the experiments using live zebrafish were approved by the Committee for Animal Care and Use of Saitama University and conducted under the regulations.

### Method details

#### Genotyping

Genotyping of the mutants used in this study was determined by PCR as previously described. Briefly, the genomic DNA was extracted by NaOH method from the caudal fin of live fish, fixed larvae, or post-stained embryos, depending on the situation. Using the genomic DNA as a template, PCR was performed as described previously for the genotyping of *hox* cluster-deficient mutants (Yamada et al., 2021). After the reactions, the PCR products were separated by electrophoresis in 2 % agarose gel in 0.5 x TBE buffer.

#### Alcian blue staining of jaw cartilage in zebrafish

Zebrafish larvae at 5 dpf were fixed with 4 % paraformaldehyde in PBS. To facilitate the extraction of genomic DNA required for genotyping, the tail region was dissected using a disposable scalpel (Feather) under a stereomicroscope before Alcian blue staining. Genomic DNA was extracted from the dissected tail and used for PCR-based genotyping as described above. Subsequently, Alcian blue staining was carried out as previously described for the tailless larvae with the desired genotype. After staining, the larvae were mounted on a glass slide (Matsunami) in 70 % glycerol and photographed using a Leica M205FA stereomicroscope with a digital camera (Leica DFC350F).

#### Whole-mount *in situ* hybridization

Whole-mount *in situ* hybridization was performed as previously described (Thisse and Thisse, 2014). After *in situ* staining, the tail of the stained embryos was dissected with a disposable scalpel (Feather). Genotyping was performed using genomic DNA extracted from the dissected tail as described above. Stained embryos with the desired genotype were photographed under a Leica M205 FA stereomicroscope with a digital camera (Leica DFC350F).

#### Whole-mount immunostaining

To visualize the pharyngeal pouches in zebrafish, whole-mount immunostaining was performed by using zn-8 monoclonal antibody (Hybridoma Bank). Briefly, zebrafish embryos at 36 hpf were fixed with 4 % paraformaldehyde in PBS. After several washings with 0.5 % Triton X-100 in PBS, the specimens were reacted overnight at 4°C with zn-8 antibody (1:500 dilution) in PBS containing 10 % fetal bovine serum, 10 % DMSO, and 0.2 % Triton X-100. After several extensive washings with 0.5 % Triton X-100 in PBS, the specimens were further reacted overnight at 4°C with a biotin-conjugated anti-mouse IgG antibody. Then, the specimens were further reacted with Vectastain Elite ABC RP kit (VecotorLabs). A signal was obtained by incubating the samples in DAB solution.

#### Injection of antisense morpholino oligos into zebrafish embryos

Antisense morpholino oligos (MOs) were injected into fertilized zebrafish embryos, which were then incubated at 28.5 °C until the developmental stages under examination. The MOs were synthesized in Gene Tools, and their sequences are as follows: negative control MO 5’-CTCTTACCTCAGTTACAATTTATA-3’, *hoxb1a* MO 5’-GGAACTGTCCATACGCAATTAA-3’, and *hoxb1b* MO 5’-AATTCATTGTTGACTGACCAAGCAA-3’. The specificity of the *hoxb1a* and *hoxb1b* MOs has been verified in a previous study (McClintock et al., 2002).

## Acknowledgements

We thank the NBRP zebrafish for providing RW strains. The zn-8 monoclonal antibody developed by University of Oregon, Institute of Neuroscience, was obtained from the Developmental Studies Hybridoma Bank, created by the NICHD of the NIH and maintained at The University of Iowa, Department of Biology, Iowa City, IA 52242. This work was supported by KAKENHI Grants-in-Aid for Scientific Research from the Ministry of Education, Culture, Sports, Science, and Technology, Japan (23K05790 to A.K.).

## Author contributions

Conceptualization: A.K.; Investigation: S.T., N.H., J.I., T.S., R.F., M.K., Y.K., and A.K.; Writing – Original Draft, A.K.; Funding Acquisition, A.K.

---

**Table S1.** Summary of pharyngeal pouch morphology in morpholino-injected embryos. Immunohistochemical staining with zn-8 antibody was performed on 36 hpf wild-type embryos injected with antisense morpholino. Following image acquisition, the morphology of the pharyngeal pouches was assessed bilaterally in each injected embryo.

| Injection | *n* | normal | abnormal |  |
| --- | --- | --- | --- | --- |
|  |  | - | more sever | less sever |
| Control | 53 | 53 (100.0%) | 0 (0.0 %) | 0 (0.0 %) |
| *hoxb1a* MO(2ng/μl)+control MO(5ng/μl) | 40 | 36 (90.0 %) | 0 (0.0 %) | 4 (10.0 %) |
| control MO(2ng/μl)+*hoxb1b* MO(5ng/μl) | 40 | 38 (95.0 %) | 0 (0.0 %) | 2 (5.0 %) |
| *hoxb1a* MO(2ng/μl)+*hoxb1b* MO(5ng/μl) | 80 | 20 (25.0%) | 54 (67.5%) | 6 (7.5%) |

